# Focal control of non-invasive deep brain stimulation using multipolar temporal interference

**DOI:** 10.1101/2023.09.05.556444

**Authors:** Boris Botzanowski, Emma Acerbo, Sebastian Lehmann, Sarah L. Kearsley, Melanie Steiner, Esra Neufeld, Florian Missey, Lyle Muller, Viktor Jirsa, Brian D. Corneil, Adam Williamson

**Affiliations:** Institut de Neurosciences des Systèmes (INS), INSERM, UMR_1106, Aix-Marseille Université, Marseille, France; International Clinical Research Center (ICRC), St. Anne’s University Hospital, Brno, Czech Republic; Department of Physiology and Pharmacology, Western University, London, Ontario, N6A 5B7, Canada; Graduate Program in Neuroscience, Western University, London, Ontario, N6A 5B7, Canada; IT’IS Foundation for Research on Information Technologies in Society, 8004 Zurich, Switzerland; Department of Mathematics, Western University, London, Ontario, N6A 5B7, Canada; Department of Psychology, Western University, London, Ontario, N6A 5B7, Canada; Robarts Research Institute, Western University, London, Ontario, N6A 5B7, Canada; Center for Social and Affective Neuroscience, Department of Biomedical and Clinical Sciences, Linköping University, Sweden

**Keywords:** Multipolar, Temporal Interference, Focality, Stimulation, Non-Human Primate, Temporally Interfering Electric Fields

## Abstract

Temporal interference (TI) is a method of non-invasive brain stimulation using transcutaneous electrodes that allows the targeting and modulation of deeper brain structures, not normally associated with non-invasive simulation, while avoiding unwanted stimulation of shallower cortical structures. The properties of TI have been previously demonstrated, however, the problem of decoupling stimulation focality from stimulation intensity has not been addressed. In this paper, we provide a possible novel solution, multipolar TI (mTI), which allows increased independent control over both the size of the stimulated region and the stimulation intensity. The mTI method uses multiple carrier frequencies to create multiple overlapping amplitude-modulated envelopes, rather than using one envelope as in standard TI. The study presents an explanation of the concept of mTI along with experimental data gathered from Rhesus macaques and mice. We improved the focality at depth in anesthetized mice and monkeys, and using the new focality in awake monkeys, evoked targeted activity at depth in the superior colliculus. The mTI method could be an interesting and potentially useful new tool alongside other forms of non-invasive brain stimulation.

Teaser: Multipolar Temporal Interference Stimulation can produce a more focal brain stimulation at depth compared to Temporal Interference.

## Introduction

Forms of non-invasive brain stimulation have been developed for neuroscientific investigation and as promising treatments for numerous neurological diseases (schizophrenia, depression, epilepsy, etc)^1^. The primary non-invasive stimulation techniques currently are Transcranial Magnetic Stimulation (TMS)^2^ and Transcranial Electrical Stimulation (tES)^3^. Using these two methods, the applied fields typically target shallower cortical brain structures, as the strength of tES and TMS is reduced as a function of distance from the electrodes and TMS coil respectively, with both techniques reaching cortical structures with sufficient strength to effectively modulate the brain^4^. Consequently, both techniques are suited to stimulate cortical regions of the brain, however, shallower brain regions receive the frequency of stimulation at a higher value of stimulation compared to deeper brain regions, and therefore when the methods are used to stimulate deep brain regions, modulating shallower regions will be a side effect^5,6^.

In 2017, Grossman and collaborators developed a novel, non-invasive method to modulate deep brain activity focally, without activating shallower cortical regions, called Temporal Interference (TI)^7^. The method simultaneously applies two high-frequency electric fields with slightly differing frequencies, where the fields constructively and destructively interfere in time, resulting in an amplitude-modulated field. Neurons have been shown by us and by others to respond to the amplitude-modulated field^8–10^. As has been demonstrated experimentally, with careful electrode placement strategies, it is possible to restrict neuromodulation to a deep brain region where the two fields overlap without activating overlaying regions of tissue. Using standard TI with two pairs of electrodes on the scalp we demonstrated the ability to adequately activate deep targets, such as the hippocampus^9^, and to activate deep peripheral nerves, such as the sciatic and hypoglossal nerve^10,11^. However, tuning the size of the TI stimulation region and the intensity of the stimulation independently is not currently possible in a standard TI configuration using two high frequencies, but would be a valuable tool – at the moment increasing the applied amplitude of TI stimulation increases the size of the of stimulated brain region. Tuning amplitude independently could increase the utility of TI in for example brain mapping to assist identifying epileptogenic regions of an epileptic brain^12^, or potentially assist investigation of whether a patient will benefit from DBS prior to an implantation^13^. When coupled with non-invasive imaging techniques such as fMRI or MEG, a non-invasive and focal TI stimulation with independently tunable amplitude and focality would be an interesting addition to gather data in order to build tailored, patient-specific brain models, e.g. for pre-operative planning^14^

In this work, we present a non-invasive deep brain stimulation method, multipolar Temporal Interference (mTI), potentially capable of meeting the above focality and intensity challenge. We describe in detail the method of mTI and demonstrate the new technique with experimental data and its feasibility in rhesus macaques, showing non-invasive deep brain stimulation which can expand and contract the stimulation volume while maintaining the stimulation intensity. We further show that the method scales down to smaller animals by showing a significant increase in focality with mTI stimulation in mice. Finally, we apply this technique in an awake monkey to confirm the possibility to modulate electrophysiological activity in deep regions. We target the superior colliculus (SC), a midbrain node of the oculomotor system that controls gaze position. The functional contribution of this structure to saccadic eye movements and the repertoire of responses that can be evoked by intracortical microstimulation are extremely well characterized^15^. Further, the oculomotor system is ideally suited to address how activity evoked by stimulation interacts with endogenous activity present at the time of stimulation^16^. We also can assess the absence of adverse reactions to stimulation by ensuring that the animal continues to perform a behavioral task during mTI. Across two behavioral tasks, we indeed show that task performance is not influenced by mTI, and we also show for the very first time that activity in the superior colliculus can be evoked using completely non-invasive transcutaneous stimulation. The method could be useful for as an additional tool in basic and clinical applications which employ non-invasive brain stimulation for diagnostics, therapy, and brain exploration.

## Materials and Methods

### Macaque anesthetized experiment

#### Ethics and animal history

Data was obtained from a male rhesus macaque monkey (*Macaca mulatta*, weighing 15 kg), as part of a planned endpoint. All surgical and experimental procedures conformed to the policies of the Canadian Council on Animal Care on the care and use of laboratory animals and were approved by the Animal Use Subcommittee of the University of Western Ontario Council. Prior to the procedure described below, the animal had a Utah array implanted in the left prefrontal cortex which had been removed 18 months earlier, as well as a pre-existing cranial implant of dental acrylic. We relied on stereotactic coordinates^17^ and anatomical MRI and CT images to plan the placement of recording electrodes.

#### Surgical procedure and stimulation

On the day of the procedure, the animal was administered ketamine and medatomidine, anesthetized via propofol constant rate infusion and isoflurane inhalation, and positioned in a stereotactic frame. The acrylic implant was then removed. 2-mm diameter burr-hole craniotomies were performed at pre-determined sites to permit access for Stereoelectroencephalography (sEEG) recording electrodes (Alcis Depth Coagulation Electrode). Each sEEG electrode comprised 15 recording contacts, distributed over 51 mm (contact length 2 mm, inter-contact distance 3.5 mm, diameter 0.8 mm). Eight craniotomies were positioned to permit access of recording contacts within the temporal lobe. A ninth craniotomy was positioned to permit access to the left hemisphere. Following each of these nine craniotomies, the sEEG electrode was lowered through a guidance screw (Alcis 2023VG, 15-25 mm length, stainless steel) that was secured to the skull via dental acrylic as in Acerbo et al^8^. All recording contacts were connected to an Intan recording system.

Following insertion of the sEEG electrodes, 16 locations on the scalp were selected for placement of TI stimulation electrodes. Stimulation locations were placed on the skin, on the portion of the scalp around the previous acrylic implant. The scalp was shaved and cleaned with rubbing alcohol. Stimulation was delivered through standard ECG monitoring electrodes (Medi-Trace 230, Ambu, Denmark); portions of the adhesive part of each electrode were trimmed in order to fit all electrodes on the scalp. Each electrode was connected to the output of a Digitimer DS-5 stimulator, which itself was connected to a waveform generator (Keysight EDU33212A). Connections were arranged to deliver mTI (8 frequencies, creating 4 overlapping envelopes).

Contact n°3 of SEEG electrode D (Figure 1 D and E, red arrow) was located at the anatomical target – this contact was monitored during stimulation to ensure that the 4 overlapping AM signals were indeed present at the target. The current provided by each of the 8 DigiTimer DS5 stimulators (8 pairs of electrodes, creating 4 envelopes) varied between 1.1 mA and 2.6 mA, so that the measured amplitude of each AM envelope signal at contact n°3 of SEEG electrode D was equal to 750 µV peak-to-peak. The necessary variation in applied stimulation currents to achieve an identical electrical potential at brain target results from the different distances between a given pair of stimulation electrodes at the brain target.

**Figure 1.**
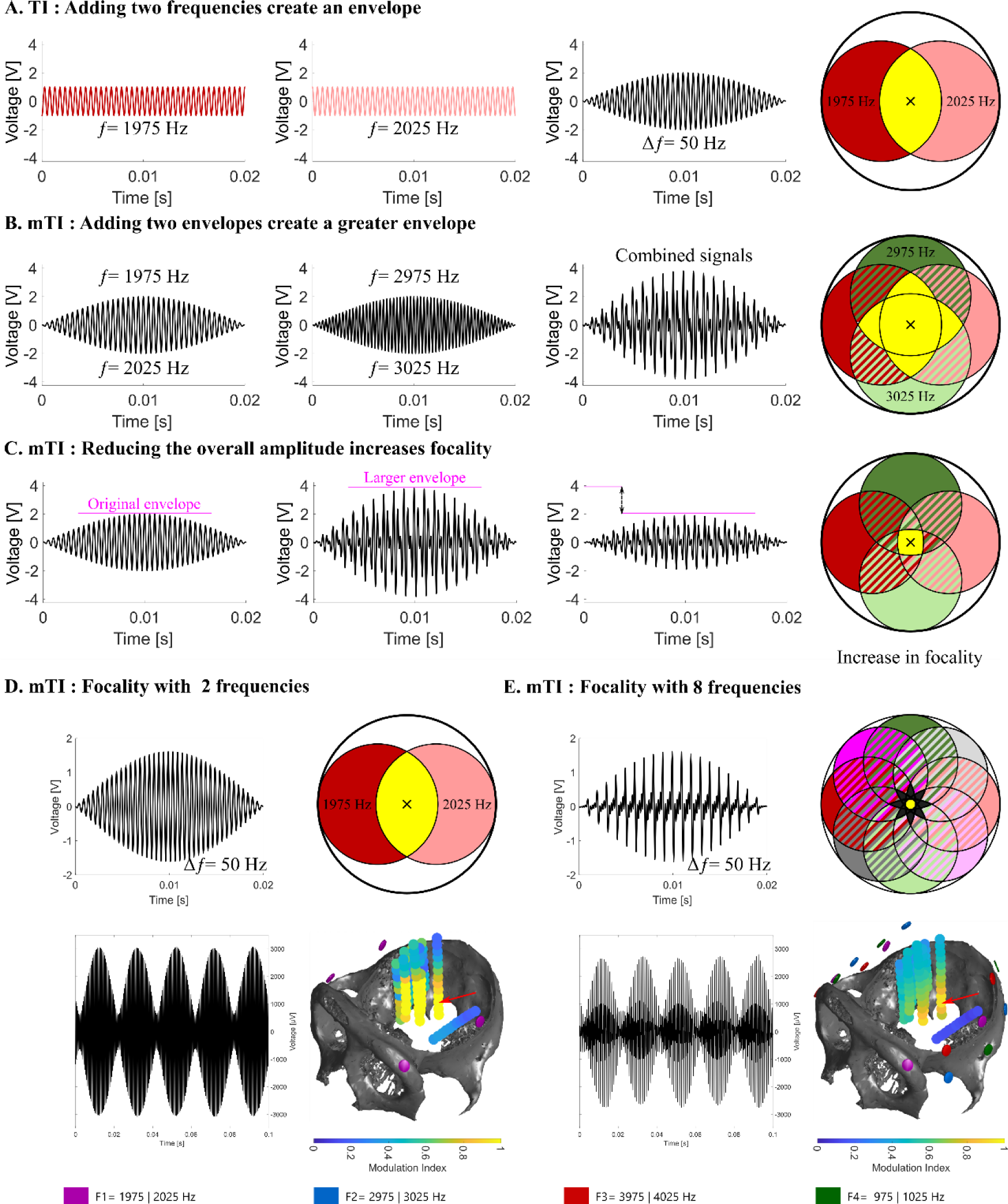
Principle of multipolar temporal interference and increased focality in non-human primates. **A. TI: Adding two frequencies to create an envelope** – Two equal amplitude sinusoidal signals at 1975 Hz and 2025 Hz interact to create a 50 Hz amplitude modulated signal. **B. mTI**: **Adding two envelopes to create a greater envelope** – The first 50 Hz envelope signal is created from the interaction of 1975 and 2025 Hz signals, the second from 2975 and 3025 Hz. When the two envelopes are added, a larger 50 Hz envelope is created. The maximum modulation occurs at the center of the circle (yellow area) **C. mTI: Reducing the amplitude to that of the original TI envelope increases focality** – The amplitude from the electrode pairs is reduced to create an envelope amplitude equal to the original standard TI envelope with only two frequencies. The focality will increase compared to the standard TI, while maintain the AM maximum of the standard TI. **D. Classic TI and measurements a macaque monkey –** 3D reconstruction of the skull of the macaque monkey (CT scan) with depth electrodes used to record the TI amplitude modulation. The region of strong amplitude modulation is found to be subcortical with TI using two high frequencies, but the focality is limited. Modulation index is calculated as follow MI=1-(min/max)), also illustrated in Fig. 2A. Position of stimulation electrodes (skin) are shown as pink circles around the skull. Red arrow shows the contact of the electrode which recorded the maximum MI. **E. mTI and measurements in a macaque monkey –** The focality of the AM signal compared to D is improved with mTI using 8 frequencies to create 4 overlapping envelopes, where the amplitude from the pairs of carrier frequencies have been reduced to create an envelope with equal amplitude to D, which results in a more focal spatial profile. As in 1D, the position of stimulation electrode (skin) are shown as pink, blue, red and green circles around the skull. Red arrow shows the electrode which recorded the maximum MI.

The applied mTI was as follows: First AM signal (1975 and 2025 Hz), Second AM signal (2975 and 3025 Hz), Third AM signal (3975 and 4025 Hz) and Fourth AM Signal (975 and 1025 Hz). As seen (see Supplementary Figure 1), it is necessary the four envelope signals be in phase, envelppe. For the experiment here, the recording electrode at the target was monitored to endure phase alignment of all 50 Hz envelopes.

#### Perfusion and CT scan

Following completion of data collection, all sEEG electrodes were cut at the top of the guide tube, and each sEEG was glued to the guide tube using medical grade silicon. The animal was then deeply anesthetized with a bolus of propofol and high percent isoflurane and euthanized through trans-cardiac perfusion with cold saline, followed by 4% PFA (performaldehyde). Care was taken during euthanasia and perfusion to ensure that the guide tubes were not disturbed, and that the sEEG electrodes remained stable within the guide tube. A post-mortem CT was performed to reconstruct the location of the sEEG electrodes, via the software MRIcroGL 1.2.20210317 x86-64.

#### Quantities of interest

The work here demonstrates a synchronous utilization of several classic TI stimulation envelopes with an AM frequency, as well as the phenomenologically observed steering behavior^7^ (shift towards the weaker current, as the channel-current ratio changes) suggest that the envelope modulation magnitude is a suitable metric for TI stimulation, as the weaker of the two fields principally defines the modulation magnitude. However, the difference frequency is actually not part of the exposure spectrum; it only results from non-linear effects that produce frequency mixing. The lowest and therefore likely the strongest contribution results from the quadratic term (the next-higher relevant term is already fourth order). The low frequency component is then extracted through low-pass filtering (potentially originating in the biophysics of the capacitive cellular membrane that dampens high-frequency responses). On that basis, a root-mean-square metric (with a running box-car average for ‘mean’) is employed to translate transient high-frequency exposure into a meaningful ‘modulation envelope’, the modulation amplitude of which is used as mTI exposure metric throughout this study, as seen in Figure 3. For the two channel case, the resulting metric is equivalent to the envelope obtained when fitting the peaks of the high-frequency signal. However, it generalizes naturally to more than two frequencies (mTI) and accounts for the varying oscillation density apparent when more than two channels are present.

#### Mock signals

Scalar signal-forms for theoretical analysis and visualization were generated by plotting simple sine functions in Matlab R2020b, such as “*sin(2*pi*frequency*t)*”. Amplitude modulated signals were obtained by adding several sine functions. Note that analyses using such scalar signals are illustrative only, as in reality, the electric fields are vectorial and the transient field evolution is more complex. However, when neural responsivity is strongly dominated by a certain field orientation (e.g., fiber orientation, orientation of pyramidal neuron axis) and the projected field component along that direction can be considered relevant, the vectorial case essentially reduces to a scalar one. The field distribution illustrations in Figure 1, were created with the vector graphics software Inkscape 1.0.2-2.

#### Signal processing

The recordings used for the modulation index (defined as MI=1-(min/max)) analysis in Figure 1 are 0.2 second samples that are representative of the recordings at calibration point D3 for the various configurations, after phase and amplitude calibration. The recordings used for the analysis in Figure 1 were digitally filtered to keep only the high carrier frequencies and to allow an analysis of the stimulation artifacts without being influenced by physiological events or noise. For that purpose, a high-pass finite impulse response filter with a stop band of 500 Hz was applied to all signals. To highlight the high frequencies carriers, three band stop filters attenuating the power of the frequencies between 1300 and 1700, 2300 and 2700, 3300 and 3700 Hz were applied to the signals. Finally, a low pass filter of type IIR with a stop band of 4300 Hz was applied to all signals. The modulation index plotted in Figure 1 D and E was calculated from the filtered signals (see above). The signals were squared to emphasize the nonlinearity phenomenon to which the neurons are subjected. A digital low-pass filter with a stop band of 200 Hz was applied to the signal, then the square root of the real part was taken. See Appendix for a discussion on this choice of the TI exposure metric. For each parameter (TI and mTI) the MI values were plotted at their respective coordinates in Figure 1 D and E. The skull was also plotted to provide spatial orientation.

#### Skull geometry

The skull geometry was reconstructed from a CT scan of the monkey with the software InVesalius 3.1.1 and was exported to Blender 3.1 where it was simplified through a decimation process. The top part of the skull was also removed with Boolean operations to provide a window for visualization.

#### Electromagnetic Field Computation

Electric field simulations were performed on the Sim4Life platform (Zurich MedTech AG) using the ‘stationary current solver. This solver is a real-valued quasi-electrostatic finite element methods (FEM) solver using the ohmic current approximation. The simulation solved the Laplace equation, ∇·σ∇Φ=0, where σ and Φ are the electric conductivity and potential respectively. The solver is suitable for the frequencies in this paper, as it has been used to model TIS stimulation in other FEM models meant to approximate the distribution of TIS that have shown accuracy.

A generic model of a rhesus macaque including over 350 different tissues was used from the ITIS ViZoo database^18^, which had been created from cryosection data. The solver discretizes the model using rectilinear meshes with nodes every 0.5 mm, spanning the entire head, down to the collar bones. This node size was chosen using a computational analysis comparing node sizes from 0.2mm to 1mm, choosing the size that minimizes computational complexity while maintaining a high degree of accuracy. The conductivities of the homogenous tissues were assigned according to the IT’IS Foundation database of tissue properties which uses Cochrane’s formula to combine standard deviation values provided by the literature. As the ohmic approximation implies, the same conductivity values were used at all frequencies. The simulations were performed with Dirichlet boundary conditions at active electrodes. The stimulating currents were normalized by integrating the current density across a plane transversing the whole model set between the electrodes.

### Mouse experiment

#### Ethics

The experiment was performed in accordance with European Council Directive EU2010/63, and French Ethics approval (Williamson, n. APAFIS#20359-2019041816357133 v10). 12-week-old OF1 mice were maintained in a transparent cage in groups of 5 mice, in a temperature-controlled room (20 ± 3°C) with a 12/12h night/day cycle. The animals had ad libitum access to food and water.

#### Surgical procedure

The mouse underwent a surgical procedure to implant minimally invasive screws to perform TI stimulation. The mouse was anesthetized via an intraperitoneal injection of a mix of Ketamine (50mg/Kg) and Xylazine (20mg/Kg) before being placed in a stereotaxic frame. After a midline incision, a verification that the lambda and bregma were on a horizontal plane was performed, before getting bregma coordinates. 8 pairs of minimally invasive electrodes were implanted around a multi-site depth probe (32 channels spaced by 100µm, NeuroNexus A1x32-Edge, USA). Each pair was placed at 1mm from the depth probe [AP: - 2.7, ML: +2.04, DV: 3.2]. A reference for the depth electrode has been placed in the right part of the cerebellum. Stimulation procedures occurred directly after implantation of stimulating and recording electrodes.

#### Electrophysiological recordings and stimulation

All electrophysiological recordings were performed with a recording/ stimulation controller (IntanTech, Intan 128ch Stimulation/Recording Controller) with a sampling rate of 30kHz. To process the data, all files have been converted from RHS format to MAT. To perform stimulation, waveform generators (Keysight, USA) were driving independent current sources (Digitimer, UK), which were stimulated via the implanted cortical electrodes. The stimulation protocol is the same as the one used in the anesthetized macaque experiment. The mTI stimulation was realized with the following frequencies: 975|1025, 1975|2025, 2975|3025 and 3975|4025 Hz. The Dipole stimulation was realized with the frequency 1975|2025 Hz. All sets of frequencies resulted in the creation of envelopes of 50 Hz. The amplitude reached by the stimulation artifact is around 3200 µV for both TI and mTI at the target contact number 16. Stimulation was continuous for each configuration.

#### Calculation of modulation Index

The modulation index shown in Figure 2 B was calculated in the same way as for the anesthetized macaque experiment in Figure 1 D-E.

**Figure 2.**
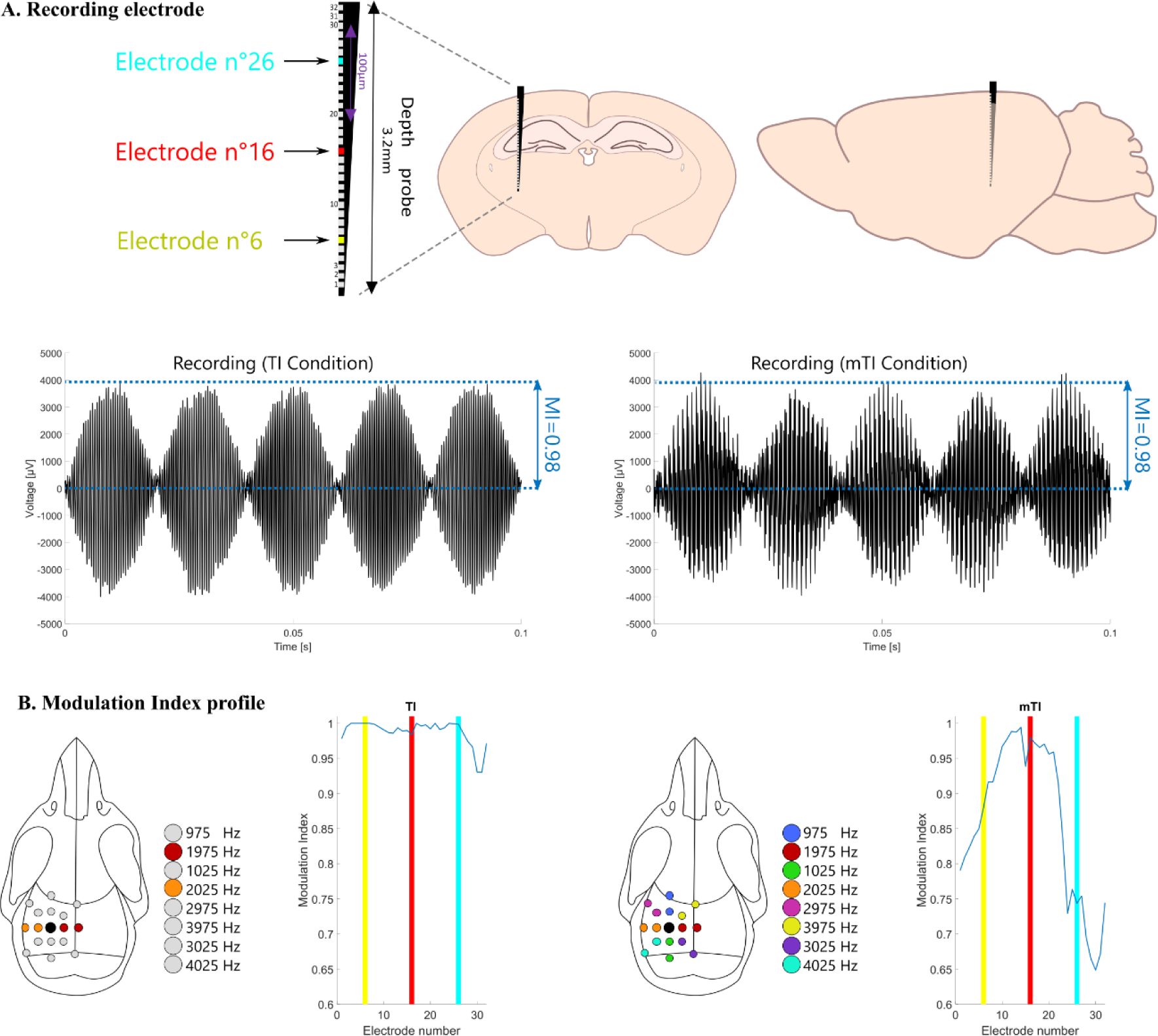
Increased Focality in rodents. **A. Experimental mouse set up** – Position of the recording electrodes in the mouse brain [AP: -2.7, ML: +2.04, DV: 3.2]. Plots of raw recording from electrode n°16 (highlighted in red) for standard TI with two frequencies and mTI with 8 frequencies. **B. Spatial distribution of Modulation Index** – Shallow (electrode 26 - blue) and deep contacts (electrode 7 - yellow) are highlighted to facilitate localization in the modulation index plots. The maximal modulation index for both TI and mTI configuration is reached in the hippocampus (approximately electrode 16 – red). However, the focality is improved in the mTI configuration. When comparing these results with those from the previous and subsequent figures, we conclude that mTI scales well between larger and smaller anatomical geometries.

#### Illustrations

The illustrations depicting the recording electrode, mouse brain, and mouse skull in Figure 2 were created using Inkscape.

### Macaque awake experiment

#### Ethics and animal history

Stimulation was also delivered to a different male rhesus macaque monkey (macaca mulatta, weighing 10.5 kg), during the performance of behavioral tasks. All training, surgical, and experimental procedures were in accordance with the Canadian Council on Animal Care policy on the use of laboratory animals and were approved by the Animal Use Subcommittee of the University of Western Ontario Council on Animal Care. We monitored the monkey’s weight daily, and their health was under the supervision of the university veterinarian. A cranial implant of dental acrylic secured a titanium headpost, as well as a recording chamber placed over a 19-mm diameter craniotomy that permitted access to the superior colliculus. This animal had a previously implanted Utah array in the left prefrontal cortex that had been disconnected over 1 year prior to the collection of the reported data. Surgical details regarding our procedures for the preparation for SC recordings in awake behaving animals have been published previously^19,20^.

#### Behavioral tasks

As depicted in Fig. 3A, the animal was placed with their head restrained 33 cm in front of a 42-in color monitor (4202L Elo Touch Solutions, Inc, Milpitas, CA) in a dimly-lit, sound-attenuated room. Eye movements were monitored using an ISCAN eye tracker which measures horizontal and vertical eye position as well as pupil dilation at 120 Hz. General behaviour was monitored remotely via a video camera. Visual stimuli and task behaviour were controlled by custom-built programs (MonkeyLogic^21^ for Matlab), which also triggered delivery of TI and SHAM during various phases of the behavioral tasks.

**Figure 3.**
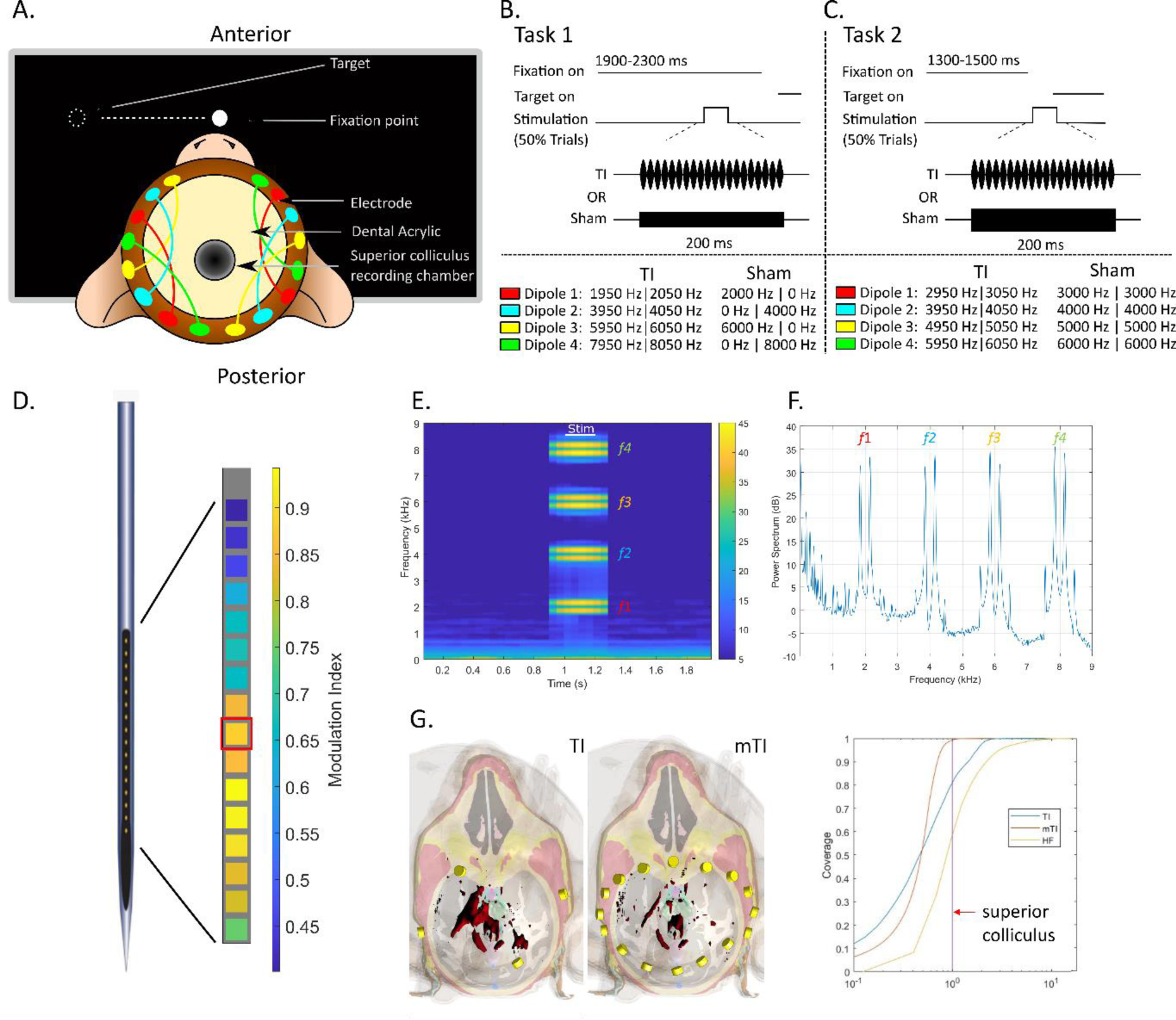
mTI layout, behavioral paradigm, and simulations in the awake behaving macaque. **A. Experimental setup.** Sketch of the placement of scalp-based stimulation electrodes around the acrylic implant in which the SC recording chamber was embedded. The stimulation electrodes are color coded and connected by colored lines to show the pairs of electrode for mTI. **B,C. Behavioral tasks.** 200 ms of mTI stimulation was embedded either within a visually-guided saccade task where stimulation was either delivered during a period of prolonged visual fixation (Task 1) or around the time of target onset in a visually-guided saccade task (Task 2). We also show the settings of the carrier frequencies for the delivery of mTI stimulation or SHAM stimulation. **D. Focusing mTI**. An intermediate contact on the linear electrode array (shown in red) located within the intermediate SC was selected to be the target for mTI. **E. Spectrogram of signal -** Frequencies used for the stimulation are seen in a time-frequency plot. For this experiment, a 100 Hz envelope was used to evoked activity. **F. PSD of signal -** Time frequency and periodogram of one of the 200ms of TI – here only mTI is displayed. The different frequencies applied are visible and a lowpass filter <100Hz is enough to remove the artefact of stimulation in the recorded data. The ability to filter stimulation artifacts is an advantage of the TI method, due to the stimulation artifact typically being several thousand Hz above the electrophysiological activity of interest. **G. Simulations of TI and mTI stimulation.** Simulation of the volume of tissue activated following either TI or mTI stimulation. (surface in red showing 6 V/m or higher). The iso-surface from mTI is more focal than that from TI. Furthermore, the cumulative histogram shows the normalized maximum electric field at the target (superior colliculus) vs the rest of the brain. For the mTI (orange), less than 1 percent of the brain receives an electric field higher than the electric field in the colliculus. For the TI (blue), more than 20 percent of the brain receives an electric field higher than the electric field in the colliculus. For context, we see that 40 percent of the brain receives a higher electric field from the high frequency carriers (yellow), which is to be expected as the carrier frequency maximum is closest to the electrodes providing the carrier.

The animal was trained on two tasks requiring them to look from a central fixation point to a peripheral visual target for liquid reward. Stimulation was delivered on half of all trials either during a period of active fixation (Task 1) or during saccade preparation (Task 2). For both tasks, the peripheral visual target was presented in the response field of the recorded SC neurons (∼12 degrees horizontal, contralateral to the side of recording). Task 1 required the animal to look steadily at the central fixation point for a prolonged period (1900-2300ms). In the 50% of trials where stimulation was delivered, 200 ms of stimulation could start 1200-1400 ms into the fixation interval (Fig. 3B). In Task 1, stimulation was delivered when neural activity in the vicinity of the recording electrode is relatively inhibited^22^, and the long fixation period facilitates measurement of pupil dilation^23,24^.

In Task 2, TI stimulation was timed so that the post-stimulation interval would overlap with visual and saccadic activity related to looking to the appearing peripheral visual target, which begins about 40 ms after target presentation^25,26^. For Task 2, we reduced the central fixation interval to between 1200-1500 ms, and introduced a 100 ms gap interval between disappearance of the central fixation period and peripheral target presentation. On trials when stimulation was delivered, the 200 ms stimulation interval was timed so that it ended approximately 30 ms after target presentation, which is just before the arrival of visual information in the SC (Fig. 3C).

#### Stimulation and recording procedures

TI stimulation was delivered from 8 stimulators via 8 electrode pairs - specifically 16 standard ECG electrodes (Medi-Trace 230, Ambu, Denmark) arranged in a ring around the existing acrylic implant on the skin (Fig. 3A), using the same preparation procedures as described above for the anesthetized macaque experiment. Electrodes were glued to the scalp using a silicon elastomer (Kwik-Sil, World Precision Instruments). Each electrode was connected to the output of either a Digitimer DS-5 stimulator connected to a waveform generator (Keysight EDU33212A) for Task 1, or a Digitimer DS-4 stimulator connected to a Tektronix AFG1022 waveform generator for Task 2. Connections were arranged to permit delivery of mTI through up to 4 TI envelopes.

Neural activity was recorded with an Intan RHS 128ch Stimulation/Recording Controller (IntanTech) system at 30kHz that also digitized at 1 kHz analog signals related to eye position, pupil dilation, and other task relevant events. To record SC activity, a 16-contact laminar electrode (Fig. 3D; S-probe, Plexon Inc, Dallas TX; 300 μm inter-electrode spacing) was lowered through a guide tube, and advanced until neural activity was encountered that was clearly related to the generation of visually-guided saccades. Since the electrode approached the SC in a surface-normal manner, functionally-related activity was seen across multiple channels that spanned the superficial and intermediate layers of the SC (Supplementary Fig. S2). The electrode contact which was clearly in the intermediate layers of the SC, the anatomical target, was chosen (seen in Figure 3D) and monitored to ensure that the mTI signal was indeed present (4 equal amplitude and in phase AM envelopes). The measured amplitude of each AM envelope was 400µV, corresponding to applied currents of between 0.5mA to 2mA for each Digitimer DS5.

During data collection, two blocks of trials were run. In the first block of trials, trials with mTI stimulation using a 100-Hz envelope were intermixed with trials without stimulation at equal rates (i.e., 50% TI stimulation and 50% no stimulation trials). In the second block of trials, SHAM TI stimulation trials were intermixed with no stimulation trials (i.e., 50% SHAM stimulation and 50% no stimulation trials). SHAM stimulation was delivered by either turning off half of the stimulators (Task 1; Fig. 3B), or by removing the frequency offset with each stimulator provided the same carrier frequency, i.e. 2000 Hz and 2100 Hz for the stimulation and 2000 Hz + 2000 Hz for the SHAM (Task 2; Fig. 3C).

Multi-unit neural spiking activity was sorted offline in principal component space after manual threshold setting (Offline Sorter, Plexon Inc., USA), and further analysed with Matlab (MathWorks Inc, USA). For firing rate visualization, spike times were convolved with an excitatory post-synaptic potential spike function^20,27^ and rasters and trial-averaged spike density functions were plotted using the Matlab toolbox Gramm^28^. Due to the artifact during the stimulation interval, we focus on spiking activity immediately following stimulation offset, as neural activity has been shown to be commonly modulated in this interval during various forms of non-invasive brain stimulation in behaving non-human primates^29,30^.

## Results

### Multipolar temporal interference (mTI) principle

Multipolar TI (mTI) creates a non-invasive stimulation with a potentially additional control of focality compared to TI and is conceptually presented in Figure 1 along with results in a rhesus macaque. As illustrated in Figure 1A, standard TI creates subcortical stimulation by a combination of two kHz frequencies applied through two pairs of electrodes, where the field from the first pair (red) and the field from the second pair (light red) overlap to create a region with strong envelope modulation (yellow). The frequency of the envelope is equal to the difference between the two kHz frequencies and is set to a value equal to a frequency from traditional deep brain stimulation to be non-invasively replicated – for example in the hippocampus 50Hz for excitation or 130Hz for inhibition in accordance with our previous publications on TI stimulation in epilepsy^13^. Unfortunately, the focality of standard TI stimulation (the size of the yellow overlap region in the schematic representation) cannot be tuned independently from the intensity of the stimulation (the maximum value of the envelope); for example, in Figure 1A, the size of the yellow region cannot simply be decreased to a four times smaller spot while maintaining the same amplitude. The problem is potentially resolved by mTI (Figure 1B).

As illustrated in Figure 1B, mTI combines TI envelopes which have different values of carrier frequency (f1 and f2 do not equal f3 and f4) but have identical envelope frequencies (|f1 - f2| = |f3 – f4| = Δf) to create overlapping regions of AM modulations. For any envelope frequency used in an mTI configuration, we have a difference of at least 1kHz between the frequencies that create a first envelope and the frequencies which create a second envelope, specifically as an example; 4000 Hz and 4010 Hz create a 10 Hz envelope, 5000 Hz and 5010 Hz create a 10 Hz envelope, however the difference between the average value of the two pairs is |4005 Hz – 5005 Hz| = 1000 Hz. The difference frequency of at least 1kHz ensures that interaction between frequencies used in the creation of a first low-frequency stimulation envelope (i.e. 10 Hz envelope) does not create additional low-frequency stimulation envelopes (i.e. |4000 Hz – 5010 Hz| = 1010 Hz). By separating pairs of carrier frequencies by at least 1kHz, we limit unwanted low-frequency envelopes in brain regions outside of the targeted brain region.

As seen in Figure 1C, when the intensity of the stimulation from the four pairs of electrodes (red, light red, green, light green) is decreased so that the peak AM envelope amplitude is equal to the AM envelope amplitude of standard TI, focality effectively increases (the yellow region is now smaller, but has the same intensity as the original yellow region).

### Demonstration of focality gain using mTI in a macaque

As seen in Figure 1D, standard TI is performed in an anesthetized macaque monkey. Two pairs of electrodes provided two frequencies. The envelope amplitude is depicted using the modulation index (the amount of amplitude modulation; again yellow is the highest). Numerous sEEG electrode recordings in the macaque show that the stimulation is subcortical, but not particularly focal. As seen in Figure 1E, mTI is performed to increase the focality compared to standard TI (8 pairs of electrodes providing 8 frequencies, creating 4 overlapping envelopes). As evident in the macaque measurements, the focality increases: the yellow region is now much smaller but has the same intensity as the original yellow region.

### Focality gain using mTI scales to smaller animals

As depicted in Figure 2A, standard TI was performed in the mouse and compared to mTI. In Figure 2B, two pairs of electrodes provided currents at two frequencies and the envelope amplitude is again depicted using the modulation index. In a mouse, a silicon depth probe was implanted down through the hippocampal target. As can be seen in Figure 2B (left panel) the 32 electrodes do not show significant variation in the modulation index across all electrodes, indicating that for a hippocampal target, at this specific amplitude of stimulation, the stimulation is not particularly focal. As further seen in Figure 2B (right panel), mTI (8 pairs of electrodes providing 8 frequencies generating 4 overlapping envelopes) increases the focality compared to standard TI, where a sharp increase can be seen at electrode 16 in the hippocampal target and decreasing above and below the target, creating a more focal stimulation for this specific AM modulation value. These results similarly show what has been shown in the previous figure, the focality significantly increases, as evidenced by the reduction of modulation index outside of the target, here a hippocampal target (around electrode 16).

### Activity evoked by multipolar temporal interference stimulation in an awake macaque

To further study the possibility of inducing an electrophysiological response, we delivered mTI stimulation to the subcortical midbrain superior colliculus (SC) in an awake macaque engaged in a prolonged period of active visual fixation (Task 1, Fig. 3B), or just before the generation of a visually-guided saccade (Task 2; Fig. 3C). Simultaneously, we recorded activity from SC via a linear array electrode positioned to span the superficial and intermediate areas of the SC. mTI stimulation was focused on an intermediate contact on the linear array from which functionally related activity was recorded (e.g., Supplementary Figure S2). For both Tasks, one block of trials consisted of mTI stimulation (8 pairs of electrodes, creating 4 envelopes) intermixed with no-stimulation trials, and another block of trials consisted of SHAM mTI stimulation intermixed with no-stimulation trials (see Fig. 3B, C for frequencies delivered). For these experiments, we created a controllable burst of envelopes of 100 Hz with four pairs of high-frequency carriers offset by a common amount (Fig. 3E,F; f1=1950|2050 Hz, f2=3950|4050, f3=5950|6050 Hz, and f4=7950|8050 Hz). Simulations (Fig. 3G) confirm the increase in focality when targeting the SC with this electrode configuration, by visualizing the amplitude of the envelope’s electric field using iso-surfaces (red) where the electric field along the surface is equal, 6V/m in these specific images. Clearly the mTI is more focal than the standard TI. Additionally, the cumulative histogram is instructive, where the electric field in the target SC is plotted with respect to the electric field in the rest of the simulated tissue. The electric field in the SC is highlighted, with less than approximately 1% of brain tissue receiving a higher electric field. In contrast, for TI, approximately 20% of brain tissue is receiving a higher electric field. The high-frequency carriers are shown for clarity, where compared to the colliculus, approximately 40% of brain tissue receives a higher electric field, which is understandable as the highest electric field from the carriers is found nearest to the stimulation electrodes.

Our choice of the SC as our subcortical target is based on its well-characterized activity patterns before visually-guided saccades^25,26^, and on the saccadic and non-saccadic effects that can be provoked by intracortical microstimulation at 100 Hz^23,32,33^. We positioned the recording electrode in the vicinity of neurons within the intermediate layers of the SC that are recruited prior to the generation of visually-guided saccades. In both Tasks, we saw no evidence that either mTI or SHAM TI distracted the animal or interfered with their ability to perform the tasks. We also saw no evidence that stimulation evoked any visible discomfort, scalp twitches, or blinking responses.

In both Tasks, we were unable to reliably resolve SC activity during the stimulation interval. Accordingly, we focus on the patterns of activity in the period immediately after stimulation offset. For the data shown from Task 1 (the active fixation task), the example neuron was clearly recruited during the generation of leftward visually-guided saccade (Supplementary Figure S2). As expected for such a neuron, we observed only a low-level of tonic activity during a period of active fixation (black traces in Fig. 4A). Notably, the activity of this SC neuron increased for ∼150 ms following mTI (red rasters and spike density functions in Fig. 4A). Such a post-stimulation increase in activity was not seen on trials with no stimulation (black rasters and spike density functions in Fig. 4A), nor on the SHAM TI trials where carrier frequency is delivered without an offset (blue rasters and spike density functions in Fig. 4A). Behaviorally, mTI stimulation did not evoke any eye movements (top rows, Fig. 4A). However, mTI stimulation did induce pupil dilation (2nd row, Fig. 4A, red traces), which is broadly consistent with stimulation of the intermediate SC that is below the level required to provoke saccades^23^. Pupil dilation, albeit to a smaller degree, was also provoked by SHAM TI stimulation (2nd row, Fig. 4A, blue traces). This latter result emphasizes the need for future controls of sensory percepts that may arise from the carrier frequencies. Specifically, we did not use a ramp in the SHAM trials which could account for the effect. Additionally, repeating similar mTI delivery in human participants would also allow for the reporting of sensations arising from stimulation.

**Figure 4.**
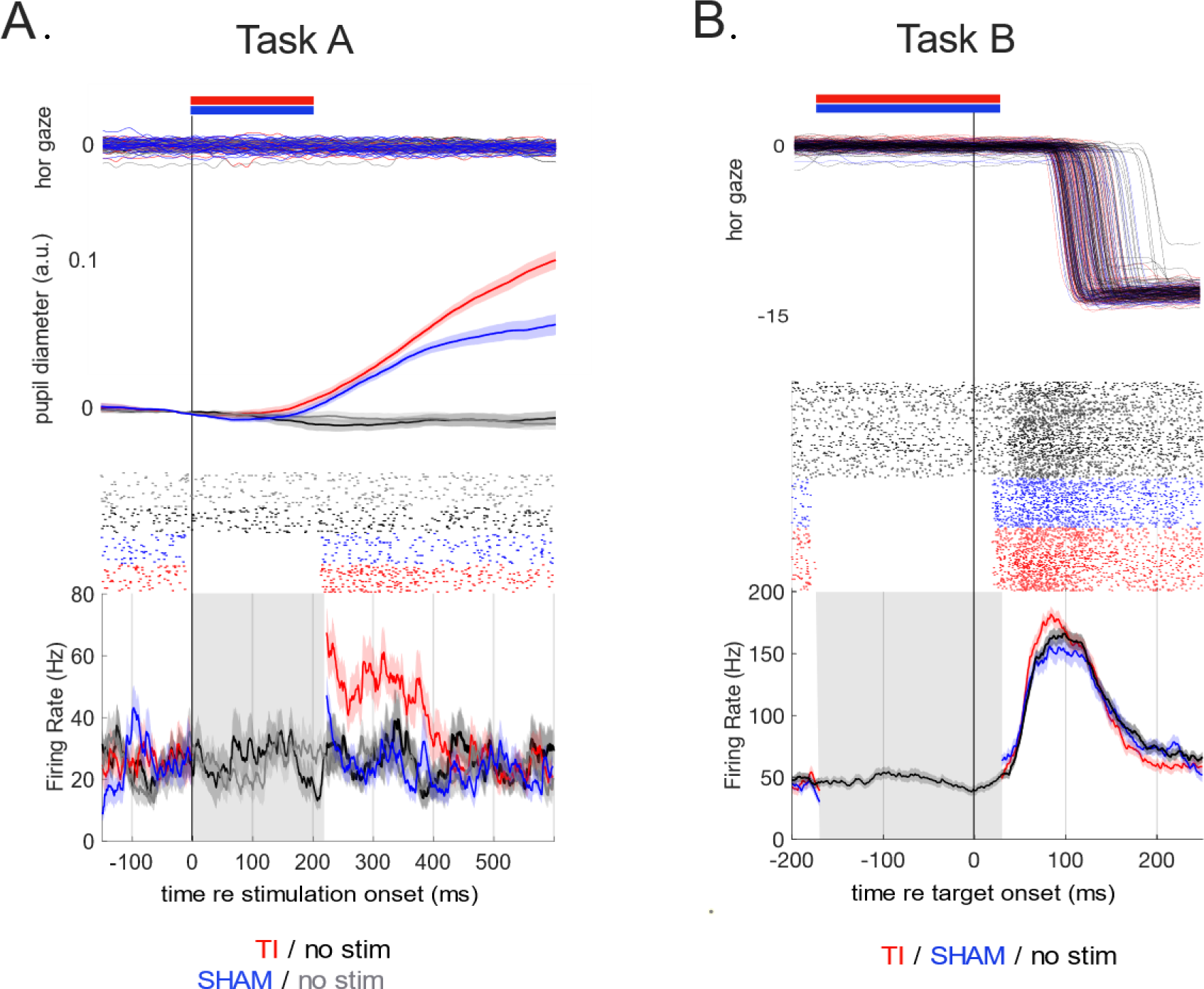
Examples of mTI on spiking activity in the superior colliculus (SC) in the awake, behaving macaque. **A) Effects of mTI during a period of stable visual fixation**. mTI did not influence eye position (horizontal gaze position, top row), but did increase SC activity (red rasters and spike density functions). Such an increase in activity was not observed on no-stimulation trials (grey and black rasters and spike density functions), nor on SHAM stimulation trials (blue rasters and spike density functions). In addition, mTI induced pupil dilation (middle row) that was absent in no-stimulaton trials and more prominent than on SHAM stimulation trials. Contours show mean subtended by the standard error of the mean. **B) Effects of mTI delivered before a visually-guided saccade.** When delivered just before the arrival of visual information in the SC in a visually-guided saccade task, mTI provoked a post-stimulation increase in functionally-related activity (red rasters and spike density functions) that was not observed on no-stimulation trials (black traces and spike density functions) nor on SHAM stimulation trials (blue traces and spike density functions).

We also studied the ability of mTI to influence functionally-related activity. On another day, our recording electrode was again positioned to record prominent saccade-related activity (see no-stimulation data shown by black rasters and spike density functions in Fig. 4B). Such functionally-related activity increased for ∼50 ms following (100 Hz envelope) mTI stimulation (red rasters and spike density functions in Fig. 4B), but did not increase following SHAM TI stimulation where all electrodes delivered high carrier frequency without any offset (blue rasters and spike density functions in Fig. 4B). We did not observe any influence of mTI on saccadic reaction time, although this may be due to the relatively low number of trials in each condition.

## Discussion

In this work, we have shown the feasibility of creating focality control with non-invasive TI brain stimulation, while maintaining the same AM stimulation amplitude. Specifically, the superposition of multiple, spatially-distinct TI-modulation regions at a single brain target was demonstrated in both non-human primates and rodents, creating a single aggregate envelope with an AM stimulation amplitude equal to the AM stimulation amplitude of standard TI however with an reduced spatial profile, improving focality (Figure 1 and 2).

The impact of TI and mTI on electrophysiological activity was further demonstrated by the post-stimulation increase in activity in the SC of an awake behaving non-human primate. To our knowledge, this is the first time that neural effects of non-invasive stimulation of the primate SC, achieved here with mTI, have been demonstrated. Given the topographic organization of the intermediate layers of the SC, future studies may be able influence specific saccade vectors by changing the focus of stimulation. mTI stimulation of the SC also provoked pupil dilation. While encouraging, we are mindful that SHAM TI stimulation also provoked pupil dilation, albeit to a lesser degree. We believe this could be related to the use of SHAM without a ramping profile. These results emphasize the need for careful behavioral controls for percepts arising from mTI in the future. Further, other work has shown that the magnitude of pupil dilation evoked by intracortical microstimulation of the SC is coordinated with subsequent saccade behavior^34^; examining whether something similar happens with mTI would also help differentiate between specific effects of mTI at the SC versus generic effects arising from sensory percepts. The feasibility of such future experiments is bolstered by our data collected to date that mTI stimulation does not cause blinking nor disrupt task performance, as could have been expected if mTI were to cause any adverse reactions. It is important to note that the sensation experienced by the macaque at the onset of the mTI stimulation was transient and typically subsides within seconds after the stimulation onset. The decision to place higher-frequency pairs closer to the animal’s face was primarily driven by practical considerations during the stimulation process.

Focality control is an interesting issue for TI^35^ where other work has set out to improve focality by adding other pairs of stimulators^36–38^. However, in these cases, the carrier frequencies creating each envelope are the same. As we have discussed in the work here, this will create a larger overlapping envelope, but the stimulation will not be more focal. As in our example, the frequencies which create a first and a second envelope, must have a difference between the average value of the two pairs which is above typical stimulation frequencies, for example 1kHz is what we have selected (i.e. |4000 Hz - 4010 Hz| = 10 Hz and |5000 Hz - 5010 Hz| = 10 Hz, however (4000 Hz + 4010 Hz) / 2 - (5000 Hz + 5010 Hz) / 2 = 1 kHz). If the difference between pairs of carriers is above 1kHz, unwanted low frequency envelope stimulation outside of the targeted brain region will be limited (i.e. |4010 Hz – 5000 Hz| = 900 Hz, there is not additional 10 Hz envelope. However, if 10 Hz envelopes are created with identical carrier frequencies, the stimulated region will be larger.

In summary, Temporal Interference (TI) represents a promising non-invasive brain stimulation technique that complements existing methods like transcranial alternating current stimulation (tACS) and transcranial magnetic stimulation (TMS). While TI has shown potential in targeting deeper brain structures without activating superficial areas, the traditional approach has been limited by the inability to decouple the intensity of stimulation from the size of the stimulated region. The work presented here introduces multipolar Temporal Interference (mTI) as an advancement in the field. By employing combinations of multiple carrier frequencies to create overlapping envelopes, mTI provides enhanced focality control, allowing for more precise targeting of deep brain regions without compromising on stimulation intensity. This novel method has demonstrated improved focality in both non-human primates and smaller animals, highlighting its potential for broader applications in both research and clinical settings.

Enhanced control over stimulation parameters could open more effective non-invasive treatments in neurological conditions, offering improved customizable and patient-specific approaches. As such, mTI not only advances the capabilities of TI but is a step to open new avenues for non-invasive brain stimulation technologies in diagnostics, therapy, and fundamental brain research.

## Acknowledgments

The authors acknowledge the generous support from the Canadian Institutes of Health Research, Canada First Research Excellence Fund (BrainsCAN), Western University, Natural Sciences and Engineering Research Council of Canada, Ontario Government, Parkinson Society Southwestern Ontario, MITACS, and the European Research Council under the Horizon 2020 program.

## Competing interest

The authors declare no competing financial interest.

All data needed to evaluate the conclusions in the paper are present in the paper and/or the Supplementary Materials.

---

**Supplementary Figure 1.**
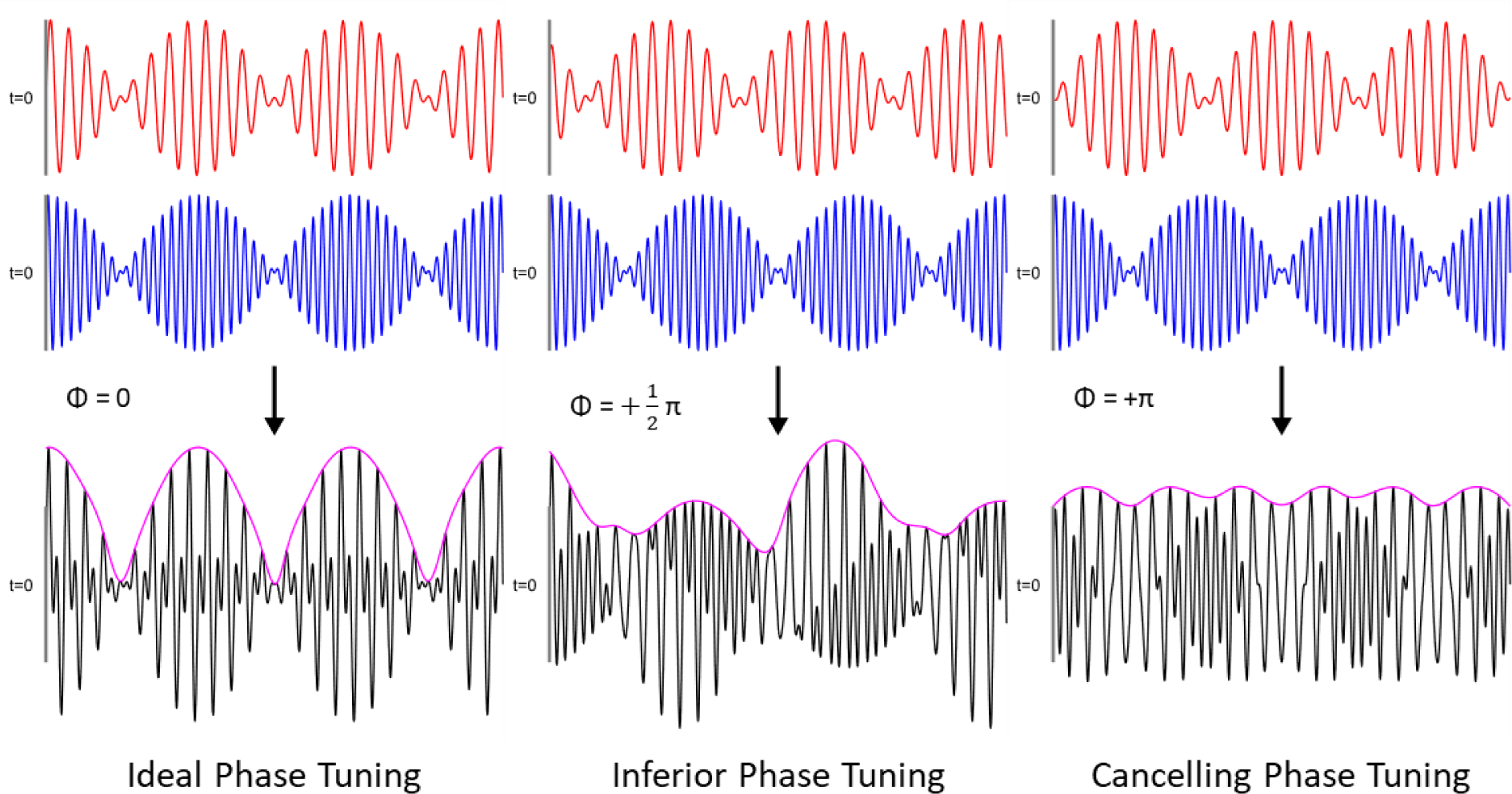
Phase interactions in mTI. (left panel) Two envelopes, in phase, create a larger aggregate envelope. (middle panel) Two envelopes, with offset phase, create an poorly defined aggregate envelope. (right panel) Two envelopes, with 180 degree phase offset, create no aggregate envelope. It is important to ensure phase alignment at the deep brain target. In this study, we ensured phase alignment as the aggregate envelope signal could be visually confirmed to be in phase using the recording electrode at the brain target. In future work, if no recording electrode is present at the target, a strategy to apply stimulation which provides in phase envelopes at the deep brain target is needed.

**Supplementary Figure 2:**
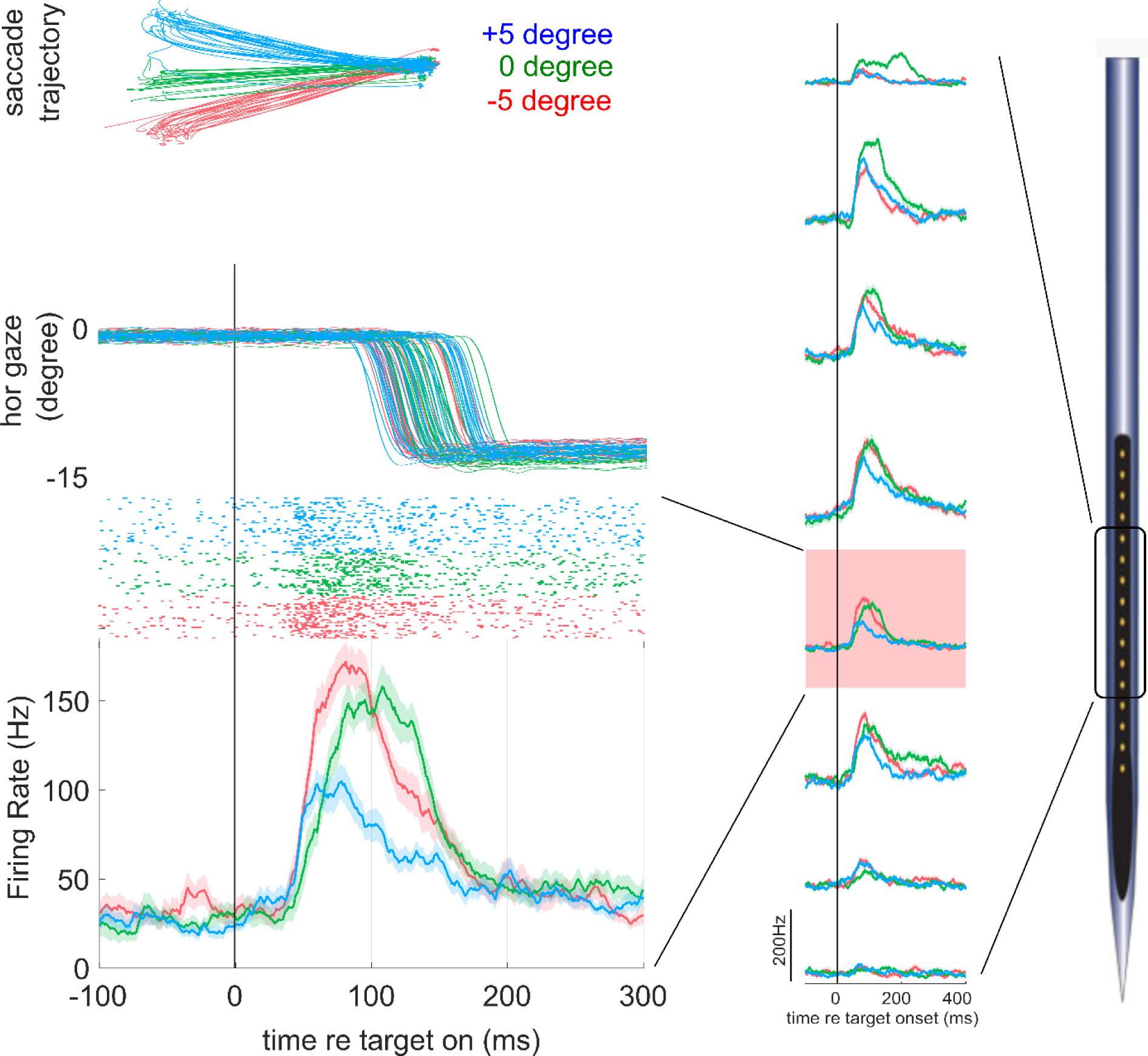
Example of functionally-related neural activity recorded from the SC. Spiking activity of the unit shown in Fig. 4A of the main manuscript, focusing here on the activity following visual target onset. Top left panel shows eye movement trajectories for three 12 deg saccade vectors directed contralateral to the side of SC recording, with the color scheme denoting different vertical components. Middle to bottom left panels show, using the same color scheme, the horizontal eye position traces (downward deflections denote left movements), the rasters of neural activity recorded from a channel in the intermediate layers of the SC, highlighted in red in the middle column, and the associated spike density functions. Middle column shows spike density functions for those channels positioned within the intermediate layers of the SC.

## References

1. Wagner T, Valero-Cabre A, Pascual-Leone A. Noninvasive human brain stimulation. Annu Rev Biomed Eng. 2007;9:527–565. doi:10.1146/annurev.bioeng.9.061206.133100

2. Klomjai W, Katz R, Lackmy-Vallée A. Basic principles of transcranial magnetic stimulation (TMS) and repetitive TMS (rTMS). Annals of Physical and Rehabilitation Medicine. 2015;58(4):208–213. doi:10.1016/j.rehab.2015.05.005

3. Roy AV, Camchong J, Lim KO. Principles and Applications of Transcranial Electrical Stimulation. In: Engineering in Medicine. Elsevier; 2019:319–334. doi:10.1016/B978-0-12-813068-1.00012-9

4. Rossini PM, Burke D, Chen R, et al. Non-invasive electrical and magnetic stimulation of the brain, spinal cord, roots and peripheral nerves: Basic principles and procedures for routine clinical and research application. An updated report from an I.F.C.N. Committee. Clin Neurophysiol. 2015;126(6):1071–1107. doi:10.1016/j.clinph.2015.02.001

5. Deng ZD, Lisanby SH, Peterchev AV. Electric field depth-focality tradeoff in transcranial magnetic stimulation: simulation comparison of 50 coil designs. Brain Stimul. 2013;6(1):1–13. doi:10.1016/j.brs.2012.02.005

6. Mikkonen M, Laakso I, Tanaka S, Hirata A. Cost of focality in TDCS: Interindividual variability in electric fields. Brain Stimulation. 2020;13(1):117–124. doi:10.1016/j.brs.2019.09.017

7. Grossman N, Bono D, Dedic N, et al. Noninvasive Deep Brain Stimulation via Temporally Interfering Electric Fields. Cell. 2017;169(6):1029–1041.e16. doi:10.1016/j.cell.2017.05.024

8. Acerbo E, Jegou A, Luff C, et al. Focal non-invasive deep-brain stimulation with temporal interference for the suppression of epileptic biomarkers. Front Neurosci. 2022;16:945221. doi:10.3389/fnins.2022.945221

9. Florian M, Rusina E, Acerbo E, et al. Orientation of Temporal Interference for Non-invasive Deep Brain Stimulation in Epilepsy. Frontiers in Neuroscience. 2021;15:633988. doi:10.3389/fnins.2021.633988

10. Botzanowski B, Donahue MJ, Ejneby MS, et al. Noninvasive Stimulation of Peripheral Nerves using Temporally-Interfering Electrical Fields. Advanced Healthcare Materials. 2022;11(17):2200075. doi:10.1002/adhm.202200075

11. Missey F, Ejneby MS, Ngom I, et al. Obstructive sleep apnea improves with non-invasive hypoglossal nerve stimulation using temporal interference. Bioelectron Med. 2023;9(1):18. doi:10.1186/s42234-023-00120-7

12. George DD, Ojemann SG, Drees C, Thompson JA. Stimulation Mapping Using Stereoelectroencephalography: Current and Future Directions. Front Neurol. 2020;11:320. doi:10.3389/fneur.2020.00320

13. Acerbo E, Jegou A, Luff C, et al. Focal non-invasive deep-brain stimulation with temporal interference for the suppression of epileptic biomarkers. Frontiers in Neuroscience. 2022;16. Accessed September 7, 2022. https://www.frontiersin.org/articles/10.3389/fnins.2022.945221

14. Ritter P, Schirner M, McIntosh AR, Jirsa VK. The virtual brain integrates computational modeling and multimodal neuroimaging. Brain Connect. 2013;3(2):121–145. doi:10.1089/brain.2012.0120

15. Gandhi NJ, Katnani HA. Motor functions of the superior colliculus. Annu Rev Neurosci. 2011;34:205–231. doi:10.1146/annurev-neuro-061010-113728

16. Lehmann SJ, Corneil BD. Completing the puzzle: Why studies in non-human primates are needed to better understand the effects of non-invasive brain stimulation. Neurosci Biobehav Rev. 2022;132:1074–1085. doi:10.1016/j.neubiorev.2021.10.040

17. Paxinos G, Huang XF, Toga A. The Rhesus Monkey Brain in Stereotaxic Coordinates. Faculty of Health and Behavioural Sciences - Papers (Archive). Published online January 1, 2000. https://ro.uow.edu.au/hbspapers/3613

18. IT’IS Database for Thermal and Electromagnetic Parameters of Biological Tissues – ScienceOpen. Accessed August 20, 2024. https://www.scienceopen.com/document?vid=a95fbaa4-efd8-429a-a59e-5e208fea2e45

19. Rezvani S, Corneil Bd. Recruitment of a head-turning synergy by low-frequency activity in the primate superior colliculus. Journal of neurophysiology. 2008;100(1). doi:10.1152/jn.90223.2008

20. Peel TR, Dash S, Lomber SG, Corneil BD. Frontal Eye Field Inactivation Diminishes Superior Colliculus Activity, But Delayed Saccadic Accumulation Governs Reaction Time Increases. J Neurosci. 2017;37(48):11715–11730. doi:10.1523/JNEUROSCI.2664-17.2017

21. Hwang J, Mitz AR, Murray EA. NIMH MonkeyLogic: Behavioral control and data acquisition in MATLAB. J Neurosci Methods. 2019;323:13–21. doi:10.1016/j.jneumeth.2019.05.002

22. Munoz DP, Fecteau JH. Vying for dominance: dynamic interactions control visual fixation and saccadic initiation in the superior colliculus. Prog Brain Res. 2002;140:3–19. doi:10.1016/S0079-6123(02)40039-8

23. Wang CA, Boehnke SE, White BJ, Munoz DP. Microstimulation of the monkey superior colliculus induces pupil dilation without evoking saccades. J Neurosci. 2012;32(11):3629–3636. doi:10.1523/JNEUROSCI.5512-11.2012

24. Lehmann SJ, Corneil BD. Transient Pupil Dilation after Subsaccadic Microstimulation of Primate Frontal Eye Fields. J Neurosci. 2016;36(13):3765–3776. doi:10.1523/JNEUROSCI.4264-15.2016

25. Goldberg ME, Wurtz RH. Activity of superior colliculus in behaving monkey. I. Visual receptive fields of single neurons. J Neurophysiol. 1972;35(4):542–559. doi:10.1152/jn.1972.35.4.542

26. Wurtz RH, Goldberg ME. Activity of superior colliculus in behaving monkey. 3. Cells discharging before eye movements. J Neurophysiol. 1972;35(4):575-586. doi:10.1152/jn.1972.35.4.575

27. Thompson KG, Hanes DP, Bichot NP, Schall JD. Perceptual and motor processing stages identified in the activity of macaque frontal eye field neurons during visual search. J Neurophysiol. 1996;76(6):4040–4055. doi:10.1152/jn.1996.76.6.4040

28. Morel P. Gramm: grammar of graphics plotting in Matlab. Journal of Open Source Software. 2018;3(23):568. doi:10.21105/joss.00568

29. Mueller JK, Grigsby EM, Prevosto V, et al. Simultaneous transcranial magnetic stimulation and single-neuron recording in alert non-human primates. Nat Neurosci. 2014;17(8):1130–1136. doi:10.1038/nn.3751

30. Romero MC, Davare M, Armendariz M, Janssen P. Neural effects of transcranial magnetic stimulation at the single-cell level. Nat Commun. 2019;10(1):2642. doi:10.1038/s41467-019-10638-7

31. Poni R, Neufeld E, Capstick M, Bodis S, Kuster N. Rapid SAR optimization for hyperthermic oncology: combining multi-goal optimization and time-multiplexed steering for hotspot suppression. Int J Hyperthermia. 2022;39(1):758–771. doi:10.1080/02656736.2022.2080284

32. Stanford TR, Freedman EG, Sparks DL. Site and parameters of microstimulation: evidence for independent effects on the properties of saccades evoked from the primate superior colliculus. J Neurophysiol. 1996;76(5):3360–3381. doi:10.1152/jn.1996.76.5.3360

33. Corneil BD, Olivier E, Munoz DP. Neck muscle responses to stimulation of monkey superior colliculus. I. Topography and manipulation of stimulation parameters. J Neurophysiol. 2002;88(4):1980–1999. doi:10.1152/jn.2002.88.4.1980

34. Wang CA, Munoz DP. Coordination of Pupil and Saccade Responses by the Superior Colliculus. J Cogn Neurosci. 2021;33(5):919–932. doi:10.1162/jocn_a_01688

35. Huang Y, Parra LC. Can transcranial electric stimulation with multiple electrodes reach deep targets? Brain Stimulation. 2019;12(1):30–40. doi:10.1016/j.brs.2018.09.010

36. Zhu X, Li Y, Zheng L, et al. Multi-Point Temporal Interference Stimulation by Using Each Electrode to Carry Different Frequency Currents. IEEE Access. 2019;7:168839–168848. doi:10.1109/ACCESS.2019.2947857

37. Cao J, Grover P. STIMULUS: Noninvasive Dynamic Patterns of Neurostimulation Using Spatio-Temporal Interference. IEEE Trans Biomed Eng. 2020;67(3):726–737. doi:10.1109/TBME.2019.2919912

38. Lee S, Park J, Choi DS, Lee C, Im CH. Multipair transcranial temporal interference stimulation for improved focalized stimulation of deep brain regions: A simulation study. Comput Biol Med. 2022;143:105337. doi:10.1016/j.compbiomed.2022.105337

