## Supplementary for "Focal control of non-invasive deep brain stimulation using multipolar temporal interference"

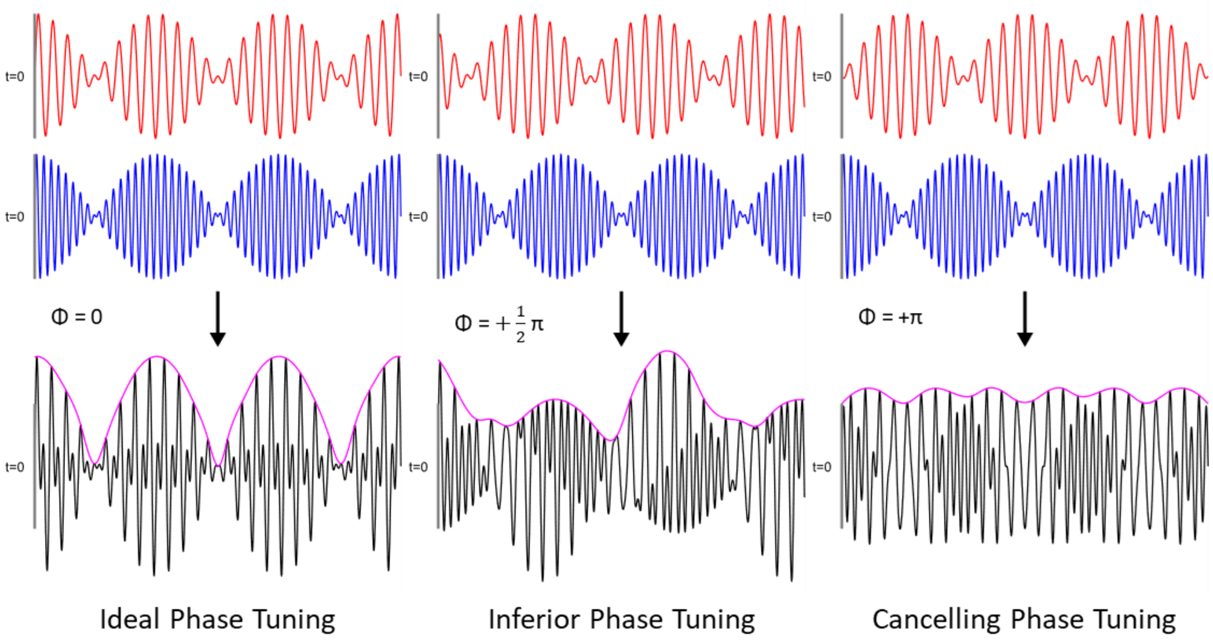


**Supplementary Figure 1. Phase interactions in a multipole**

Just like the two high-frequency carriers ('Dipole') interfere temporally in classic TI to create a low-frequency modulation envelope, multiple such Dipoles with strongly different carrier frequencies interfere to create a more complex envelope shape. However, while the envelope shape is unaffected by the relative phase in the two-channel case (only its phase is), the envelope shape has a complex dependence on the underlying phases when more than two frequencies are present. Special care must therefore be taken to ensure that the multiple Dipoles are in phase at the calibration point. The different plots here represent a case of perfect phase tuning (left), a case where the two envelopes are phased such that their combination is hardly modulated (right) and a case in between (middle).

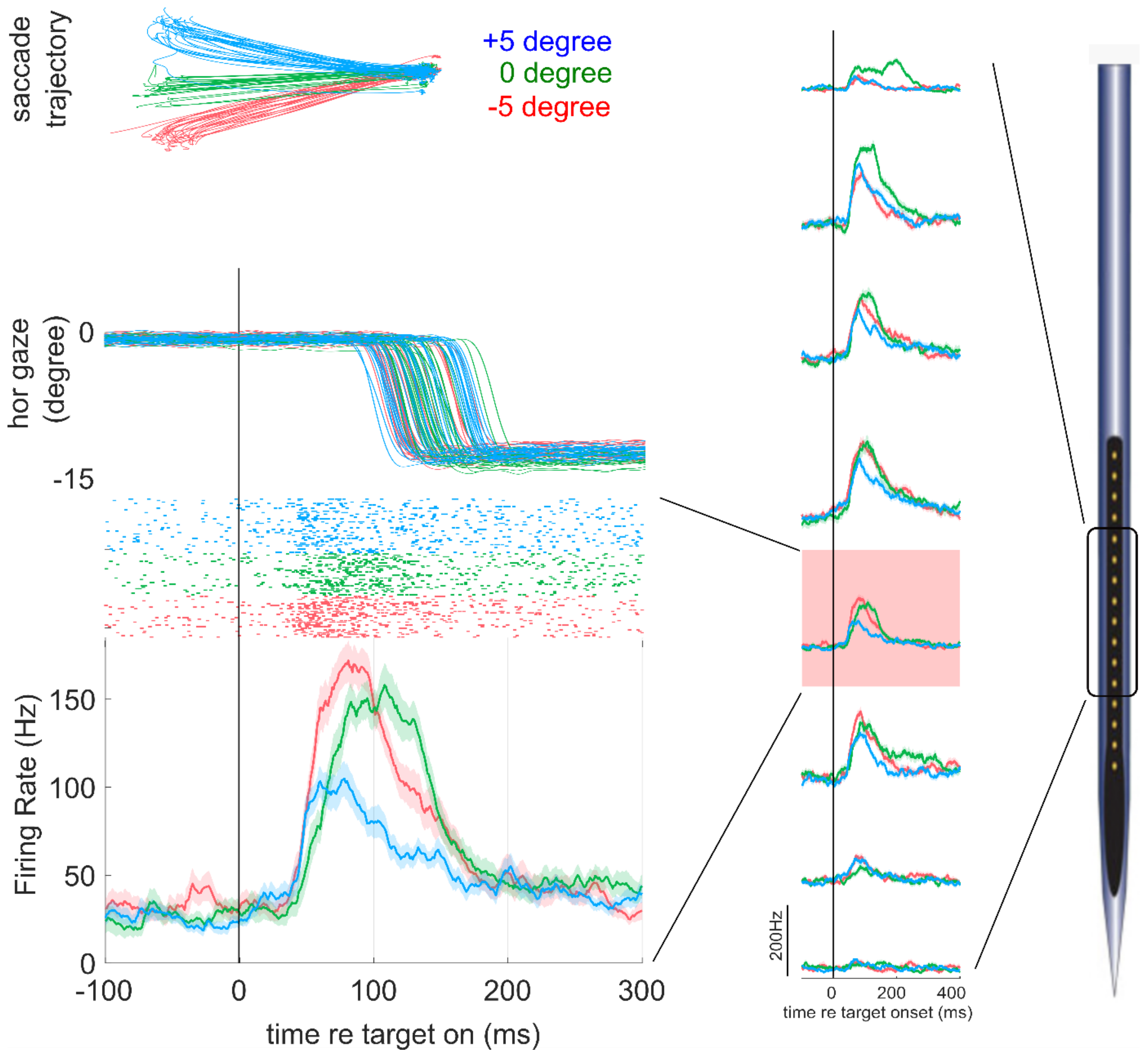


**Supplementary Figure 2: Example of functionally-related neural activity recorded from the SC** Spiking activity of the unit shown in Fig. 4A of the main manuscript, focusing here on the activity following visual target onset. Top left panel shows eye movement trajectories for three 12 deg saccade vectors directed contralateral to the side of SC recording, with the color scheme denoting different vertical components. Middle to bottom left panels show, using the same color scheme, the horizontal eye position traces (downward deflections denote left movements), the rasters of neural activity recorded from a channel in the intermediate layers of the SC, highlighted in red in the middle column, and the associated spike density functions. Middle column shows spike density functions for those channels positioned within the intermediate layers of the SC.
